# Label-free Quantification of Host-Cell Protein Impurity in a Recombinant Hemoglobin Reference Material

**DOI:** 10.1101/2023.04.14.536846

**Authors:** André Henrion, Cristian Arsene, Maik Liebl, Gavin O’Connor

**Affiliations:** Physikalisch-Technische Bundesanstalt (PTB), Braunschweig und Berlin, D-38116 Braunschweig, Germany

## Abstract

*Quantitativ*e analysis depends on pure-substance primary calibrators with known mass fractions of impurity. Here, label-free quantification (LFQ) is being evaluated as a readily available, reliable method for determining the mass fraction of host-cell proteins (HCPs) in bioengineered proteins. For example, hemoglobin-A2 (HbA_2_) is being used as obtained through overexpression in *E.coli.* Two different materials had been produced: natural, and U-^15^N-labeled HbA_2_. For quantification of impurity, precursorion (MSl-) intensities were integrated over all *E.coli* -proteins identified, and divided by the intensities obtained for HbA_2_. This ratio was calibrated against the corresponding results for *E.coli*-cell lysate, which had been spiked at known mass-ratios to pure HbA_2_. To demonstrate the universal applicability of LFQ, further proteomes (yeast and human K562) were then alternatively used for calibration and found to produce comparable results. Valid results could also be obtained when the complexity of the calibrator is reduced to a mix of nine proteins, and a minimum of five proteins is estimated to be sufficient to keep the sampling error below 15%. For the studied materials, HbA_2_-mass fractions of 916±15 mg/g and 922±11 mg/g were found. Value assignment by LFQ thus contributes 1-2% to the overall uncertainty of HbA_2_-quantification when these materials are used as calibrators. Further purification of the natural HbA_2_ yielded 999.1± 0.15 mg/g, corresponding to ≈ 0.2% of uncertainty contribution, though at a significant loss of material. If an overall-uncertainty of 5% is acceptable for protein-quantification, working with the original materials would definitely be viable, therefore.

---

Protein quantification by mass spectrometry (MS) is considered to make an essential contribution within strategies toward precision diagnostics. ^1^ Basically, uncertainties of 5%, or less, can be achieved with proteins if isotope labeled internal standards are employed (D-MS). ^2^ However, a lack of information about the impurity fraction in the calibrator-material increases the overall uncertainty and may void out the precision of results. Methods and approaches have just recently been reviewed for impurity-determination in organic compounds to be used as primary calibrators in quantitative analysis. ^3^

Rather than looking at small organic molecules, the present work is motivated by the additional need for well characterized reference materials (RMs) in targeted quantification of proteins. Depending on the measurement strategy involved, either proteotypic peptides or proteins in full-length are used for calibration. ^4-6^ For peptides, different approaches to impurity-measurement have been studied, as was reviewed in ref. 7. Direct quantification by amino acid analysis (AAA), quantitative nuclear resonance spectroscopy (qNMR), or elemental analysis were found to work best in many situations. For most accurate results, rigorous detection, quantification and correction for interfering compounds had to be involved. ^8-10^ A complementary approach consists of the one-by-one detection, identification and quantification of individual contaminants as separate analytes to obtain the mass fraction of impurity. Such mass-balance approach, in spite of being labor-intensive, is viable and common an option for short peptides. Typically, solid phase synthesis (SPSS) is used for their production. Main channels of aberration from the intended amino acid sequence are well known for SPSS.^2^ In such a setting, therefore, the number of contaminants to be taken into account may be small and manageable.

In contrast to this, practicability of both mentioned approaches is complicated for purity determination of protein materials, if not impossible at all. Indeed, effective methods are available for removing host-cell related proteins (HCPs) from the target, after expression. Still, there are circumstances that may cause significant amounts of HCPs to remain in the product. For the example of *E*.*coli, this has* been pointed out to typically happen if the expression yield is low. ^13^ Overexpression of the target might also induce expression of a number of bacterial proteins due to pleiotropism and/or stress conditions. There are recurring basic patterns of such proteins, as systematized in ref. l3. These are confined to a much smaller subset compared to the original proteome. Although this reduces their number, presence and individual abundances of HCPs may vary between preparations. At many events, residing HCPs will still be many in number, thus limiting the practicability of the one-by-one approach.

Here, we systematically evaluate the use of label-free quantification (LFQ) of proteins by precursor-ion (MSl) intensities to reliably obtain the mass fraction of HCPs in a given sample. The assumption with LFQ is, that an amount of any peptide produces a specific amount of MSl-intensity per mass of protein, regardless of what the individual peptides (proteins) are. Unlike in most applications of LFQ, ^5,6,14^ in our context, peptide intensities are not collected separately per protein, but are rather integrated over all proteins identified from the host-cell proteome. We demonstrate that this quantity can be calibrated against known amounts of the host-cell proteome. Beyond this it will be shown that, any other proteome, or even a protein mixture of just a few common proteins could likewise be used for calibration. This is being exemplified here for hemoglobin-A2 (HbA_2_), a pure-substance RM obtained by overexpression in *E.coli*. The material was produced as a primary calibrator for blood measurement of HbA_2_ by isotope dilution mass spectrometry (ID-MS). ^15,16^ Besides the natural form, it also had been engineered as U-^15^N-labeled version, thus providing an internal standard. In either case, purification by immobilized metal affinity chromatography (IMAC) had left an estimated 5−10% mass fraction of *E.coli* -proteins. In addition to the (natural) HbA_2_-RM and the (labeled) U-^15^N-RM, a third material was included in the study, that was obtained through further purification of the natural one (*ultra*-purified HbA_2_-RM). This was then used to demonstrate the applicability of the approach to low-level impurity materials as well.

## EXPERIMENTAL SECTION

### Study materials

Recombinant HbA_2_ (*α*_2_*δ*_2_) in natural and U-^15^N-labeled form was obtained from Trenzyme GmbH as previously described ^16^ using P69905 and P02042 (UniPro-tKB) as templates for co-expression in *E*.*coli* of the *α*-and *δ*-subunits, respectively. Both materials were obtained as solutions of about 0.42 mg/g (natural form) and 0.47 mg/g (labeled form) in 50 mmol/L 2-Amino-2-(hydroxymethyl)propane-1,-diol (Tris), pH 7.5, and 100 mmol/L NaCl. Recombinant HbA_2_ (*α*_2_*δ*_2_) in natural form was *ultra*-purified by semipreparative strong anion exchange chromatography using a MONO-Q 4.6/100 PE column.

### Calibrators

#### HbA_2_ used for preparation of the calibrators

The material was obtained from SIGMA-Aldrich, cat. No.: H0266; lot: SLBK8749V, as a neat substance. The (protein-) purity was 99.0% using the LFQ-method as described herein. A stock solution was prepared from this material by dissolving ∼2 mg in 1 g of Tris (10 mmol/L, pH 7.8). The mass fraction of HbA_2_ in this solution was 482.7 mg/g by AAA. An aliquot of this was used as a constant component present in each of the calibrators (red in Fig. 1A).

**Figure 1:**
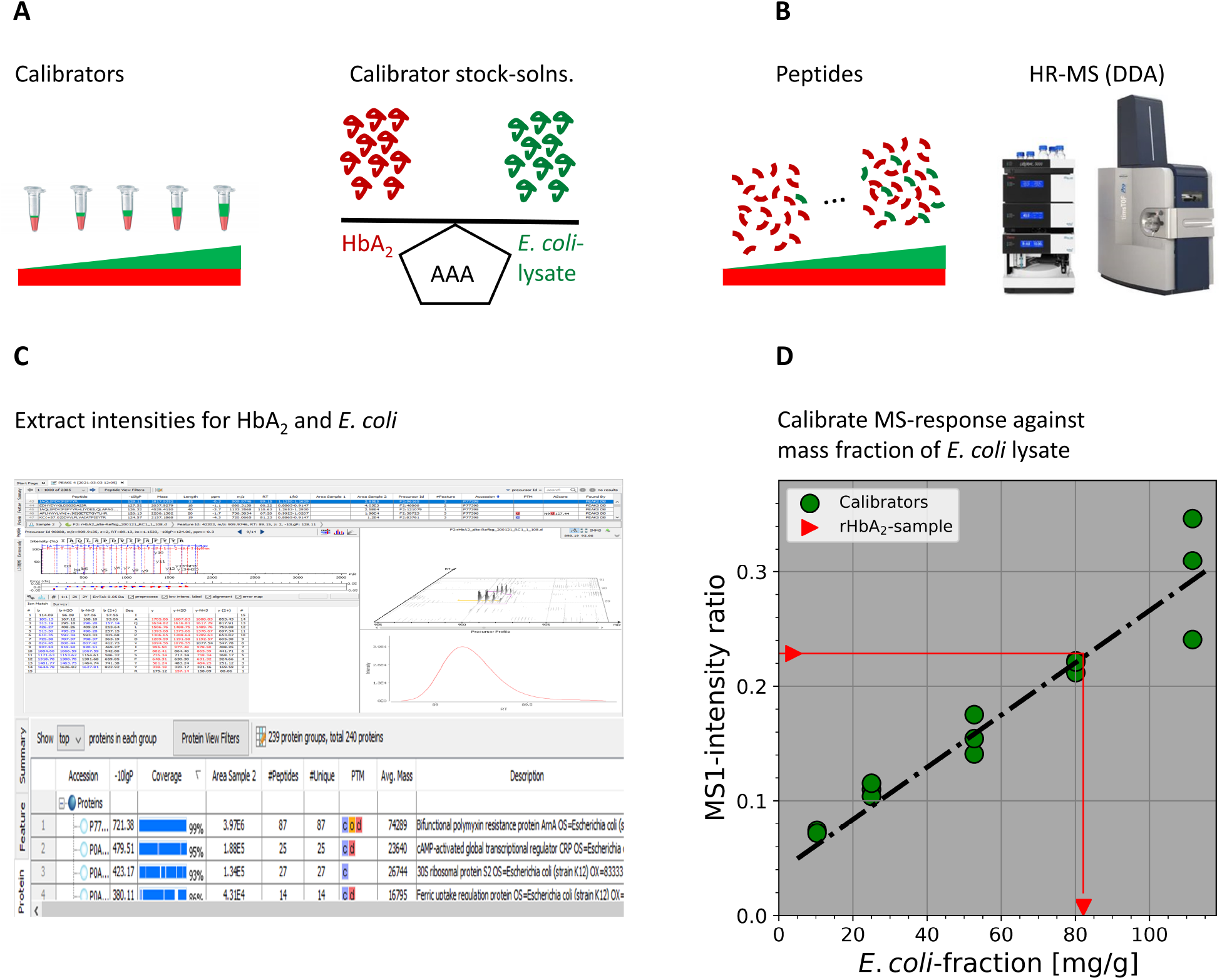
LFQ-based measurement of HCPs (*E.coli*)-mass fraction in recombinant HbA_2_. (A) Calibrators were obtained by spiking aliquots of HbA_2_-stock solution (red) with increasing amounts of *E.coli* -lysate (green). Amounts per mass (mg) of both components (HbA_2_ and *E.coli*, resp.), and mass fractions of *E.coli* were hence known for each calibrator through amino acid analysis. (B) Quantitative information was acquired by shotgun-proteomics. (C) MS1-intensities were integrated over all features associated with peptides identified from either *E.coli*, or HbA_2_. (D) The obtained ratios (*E.coli* ÷ HbA_2_) for the calibrators were plotted VS. mass fractions. The fitting linear function was then used to calculate the *E.coli* - fraction in the investigated materials from sample measurements (red line and arrows).

#### E.coli proteome-sample

Lyophiliqed *E*.*coli*-protein material was obtained from BIO-RAD (ReadyPrep^™^, Catalog 16−2110, L9703999, Control 3110004134). For a stock solution, the material was reconstituted in water (30% acetonitrile, 0.1% formic acid), mass fraction of *E*.*coli*-proteins by AAA: 0.3449 ± 0.012 mg/g. A series of calibrator samples was prepared by mixing an aliquot of HbA_2_-stock solution with the appropriate amount of *E*.*coli*-stock solution (red and green, resp. in Fig. 1A). The mass fractions of HbA_2_ in these sample solutions were: 0.390, 0.383, 0.372, 0.361 and 0.350 mg/g. The corresponding mass fractions of *E*.*coli*-protein, relative to HbA_2_ (note) in the calibrator sample were: 10.3, 25.0, 52.7, 80.1 and 111.6 mg/g.

#### Yeast proteome-sample

Yeast-based calibrators were prepared in the same way as described for *E*.*coli*. A whole-cell protein extract of *Saccharomyces cerevisiae* (Promega, V7341, lot 434786) was deployed. The material came as a solution in 50 mmol/L Tris and 6.5 mol/L urea. Mass fractions of HbA_2_: 0.390, 0.387, 0.383, 0.378 and 0.372 mg/g yeast proteins (relative to HbA_2_): 12.7, 26.5, 52.4, 76.6 and 113.6 mg/g.

#### Human K562 proteome-sample

A whole-cell protein extract from human K562-cells (Promega, V6941, lot 444583) was dissolved in 50 mmol/L Tris and 6.5 mol/L urea, as above. HbA_2_-concentrations: 0.390, 0.387, 0.383, 0.378 and 0.372 mg/g, mass fractions of K562-proteins, relative to HbA_2_: 15.0, 32.7, 63.9, 94.3 and 136.2 mg/g.

#### Protein mix

Human C-reactive protein (CRM GBW09228, National Institute of Metrology, China), human insulin analog (insulin aspart, NovoLog) and human *β*2-microglobulin (kindly provided by the Institute for Reference Materials and Measurements, Geel, Belgium) were obtained as solutions. Bovine serum albumin (SIGMA-Aldrich, cat. No. 05470, lot No. 1099572), myoglobin from horse skeletal muscle (SIGMA-Aldrich, cat. No. 70025, lot No. 381848/1), cytochrome-c from bovine heart (SIGMA-Aldrich, cat. No. C3131, lot SLBZ0555), somatotropin (NIBSC, WHO International Standard 98/574), human ceruloplasmin (Athens Research & Technology, cat. No. 16-16-030518) and serotransferrin (SIGMA-Aldrich, cat. No. T3309, lot BCBR1763V) were obtained as solids and had to be dissolved to known concentrations in water, prior to use. Mass fractions of somatotropin, ceruloplasmin, serotransferrin, *β*2-microglobulin and insulin were determined by MS based AAA, while certified values were used as provided by the supplier for C-reactive protein, albumin, cytochrome-c and myoglobin. Aliquots of these solutions were mixed to yield a stock solution containing somatotropin, ceruloplasmin, serotransferrin, *β*2-microglobulin, insulin, C-reactive protein, albumin, cytochrome-c and myoglobin in the mass-ratio of 0.1055: 0.1176: 0.124: 0.1251: 0.1247: 0.0317: 0.1215: 0.1260: 0.1233. Aliquots of this mixed solution were spiked with aliqouts of the HbA_2_ stock solution, resulting in HbA_2_-mass fractions of 0.393, 0.388, 0.383, 0.378, 0.372 and 0.367 mg/g, and protein-mass fractions, relative to HbA_2_, of 22.7, 47.4, 69.9, 92.8, 117.1 and 140.8 mg/g. Beyond this, a second series of calibrator samples was prepared, to additionally cover the low HCPs-fraction range as needed for the *ultra-purified H*bA_2_-RM. These calibrators were of 0.427 mg/g HbA_2_-mass fraction, and 0.2, 0.3, 0.4, 0.7, 1.1 and 1.4 mg/g protein-mass fractions, relative to HbA_2_.

### Determination of protein-mass fractions in the calibrators

For the stock solutions used to prepare the calibrators, mass fractions of amino acids were determined by mass spectrometry based AAA, as detailed in ref. 16. These mass fractions were then combined with the known mass fractions (or relative amounts) of these amino acids in the protein or proteome to yield the protein-mass fraction in that stock solution. In the cases of *E*.*coli*, yeast and K562, relative amounts (by mass) of amino acids were used, as was previously published (refs. 17−19). Uncertainty contributions of AAA (in our laboratory) plus uncertainties published with the literature data were combined to yield uncertainties of 3.5%, 2.7%, 3.0% and 3.2%, respectively, for the *E*.*coli*-, yeast-, K562- and protein-mix calibrator stock solutions.

### Proteolysis

To a 30 *μ*L aliquot of sample (recombinant HbA_2_) or calibrator, 70 *μ*L of Tris-solution (35 mg Tris-base, 46 mg Tris HCl, dissolved in 1 mL water) were added. Proteolysis (37 °C) was started by the addition of 10 *μ*L of trypsin solution (1 mg/mL in 50mM acetic acid). Trypsin from porcine pancreas was obtained from Sigma-Aldrich, St Louis, USA; cat. No. T0303. After 10, 70, 130, 190 and 250 min, further 10 *μ*L aliquots of trypsin solution were added. In parallel, 40 *μ*L aliquots of acetonitrile were added after 10, 30, 60, 90, 120 and 150 min. The sample or calibrator was further incubated at 37 °C overnight. For reduction, 0.8 mg of dithiothreitol (DTT) were added. After incubation (37 °C) for 1 h, 3 mg of 2-iodoacetamide were added for alkylation (30 min at room temperature). The excess of 2-iodoacetamide was quenched with 3 mg of DTT. Reaction was stopped by the addition of 10 *μ*L of formic acid 10 vol.-%). The sample or calibrator was desalted using solid-phase extraction (SPE) C18 ec cartridges (Chromabond, 100 mg, Macherey−Nagen, Düren, Germany). After lyophilization, residues were redissolved in 40 *μ*L of water 0.1 % formic acid) and subjected to nLC-MS/MS analysis.

### Liquid chromatography-mass spectrometry

An UltiMate 3000 RSLCnano HPLC system (Thermo Fisher Scientific) coupled to a timsTOF Pro mass spectrometer (Bruker Daltonik) was used for the analysis of the proteolysed sample and calibrator. Peptides were trapped on a pre-column (Acclaim PepMap C18, 5 *μ*m, 0.3×5 mm) and then separated on a Bruker Fifteen nanoFlow column (15 cm × 75 *μ*m, C18, 1.9 *μ*m, 120 Å) using a linear water-acetonitrile gradient (210 min) from 1 to 60% B and from 60% to 80% B within 20 min (with solvent A: water, 0.1 vol.-% formic acid and B: acetonitrile, 0.1 vol.-% formic acid) at 40 ° C. The flow rate was 300 nl/min. The timsTOF Pro mass spectrometer was equipped with a CaptiveSpray ion source. The mass spectrometer was run using the DDA-PASEF-standard-1.1 sec-cycletime method, as provided by Bruker. Briefly, the settings were: 10 PASEF MS/MS scans per acquisition cycle with a trapped ion mobility accumulation and elution time of 100 ms. Spectra were acquired in a *m/z* range of 100 to 1700 and in an (inverse) on mob ty range (1*/K*_0_) of 0.60 to 1.60 Vs/cm^2^. The collision energy was set-up as a linear function of ion mobility starting from 20 eV for 1*/K*_0_ of 0.6 to 59 eV for 1*/K*_0_ of 1.6.

### Protein database search

PEAKS Studio Xpro (Bioinformatics Solutions Inc.) was used for feature detection/database search and precursor ion (MS1) quantification. Databases for *E*.*coli*-, *Saccharomyces cerevisiae*-, human proteome and the mixture of nine proteins were obtained as FASTA files (uniprot.org, accessed: 27. Aug. 2021). FASTA files of human hemoglobin subunit alpha and delta (Uniprot: P69905 and P02042) were added to the databases of non-human proteomes. The following settings were applied for data analysis: carbamidomethylation of cysteine as fixed modification, methionine oxidation and glutamine or asparagine deamidation as variable modifications. A maximum of two modifications per peptide was allowed. With the human proteome, glycosylation was set as an additional variable modification using the built-in glycosylation list. Trypsin/P was set as the enzyme and no more than two missed cleavages per peptide were allowed. The tolerance for monoisotopic mass of precursor ions and fragment ions was 15 ppm and 0.05 Da, respectively. For the retention time and ion-mobility of an identified peptide the shift tolerance between different runs was 3 min and 5%, respectively. Mass correction was enabled for precursor ions. The false discovery rate (FDR) was 1% at the peptide- and protein-level. Results of quantification were obtained as peak areas at the protein-level.

### Mass fraction of impurity in labeled HbA_2_-RM

The mass fraction of impurity in labeled HbA_2_-RM was 78.1 ± 8.6 mg/g. The individual results from *n*=6 repetitions of label-free quantification were: 70.8, 72.4, 84.2, 92.8, 73.2 and 75.0 mg/g.

### Fractions of co-purifying proteins

The fraction of *E.coli* -proteins known to frequently be co-purified was calculated as the ratio of the (MS1)-intensity of these proteins to the intensity of all *E.coli* -proteins identified in HbA_2_-RM or in the *E.coli* proteome-sample. For Figure 3, fractions were calculated for the two HbA_2_-RMs and compared to the fraction for a *E.coli* proteome-sample containing a similar amount of *E.coli* proteins (80.1 mg/g).

### Downstream data analysis

Further analysis of the data exported from PEAKS was based on Python 3.8 with the modules pandas, numpy, numpy.linalg and Matplotlib imported as needed. For data-fit and cross-validation, Scikit-learn ^20^ 1.0.2 was used.

### Data availability

The mass spectrometry data have been deposited to the ProteomeX-change Consortium via the PRIDE partner repository with the dataset identifier PXD041736.

## RESULTS AND DISCUSSION

### Calibration and value-assignment to the HbA_2_-materials

The mass fractions of *E*.*coli in the HbA*_2_ (raw-) products (natural and labeled HbA_2_-RM), as well as in the *ultra-purified* HbA_2_-RM, were quantified based on a standard shotgun proteomics approach, as illustrated in Fig. l. MSl-intensity ratios (*E.coli* ÷ HbA_2_) were calibrated against known mass fractions of *E.coli* -proteins relative to HbA_2_ (Fig. lA and D). In this way, HbA_2_ technically also provides a pseudo internal standard (PIS), ^5^ improving the reproducibility of measurement. For calibration, a linear model was established for the dependence of the instrumental response (MSl intenstity ratio) on the mass fraction. In turn, as is common in quantitative chemistry, this model function was resolved for the fraction as dependent variable, thus allowing the fraction to be predicted from an intensity ratio as input (red arrows in Fig. lD). The result of such a calibration with *n* = 3 repeats at each level, and subsequent value-assignment to both HbA_2_-RMs (natural and labeled) is shown in Fig. 2A.

**Figure 2:**
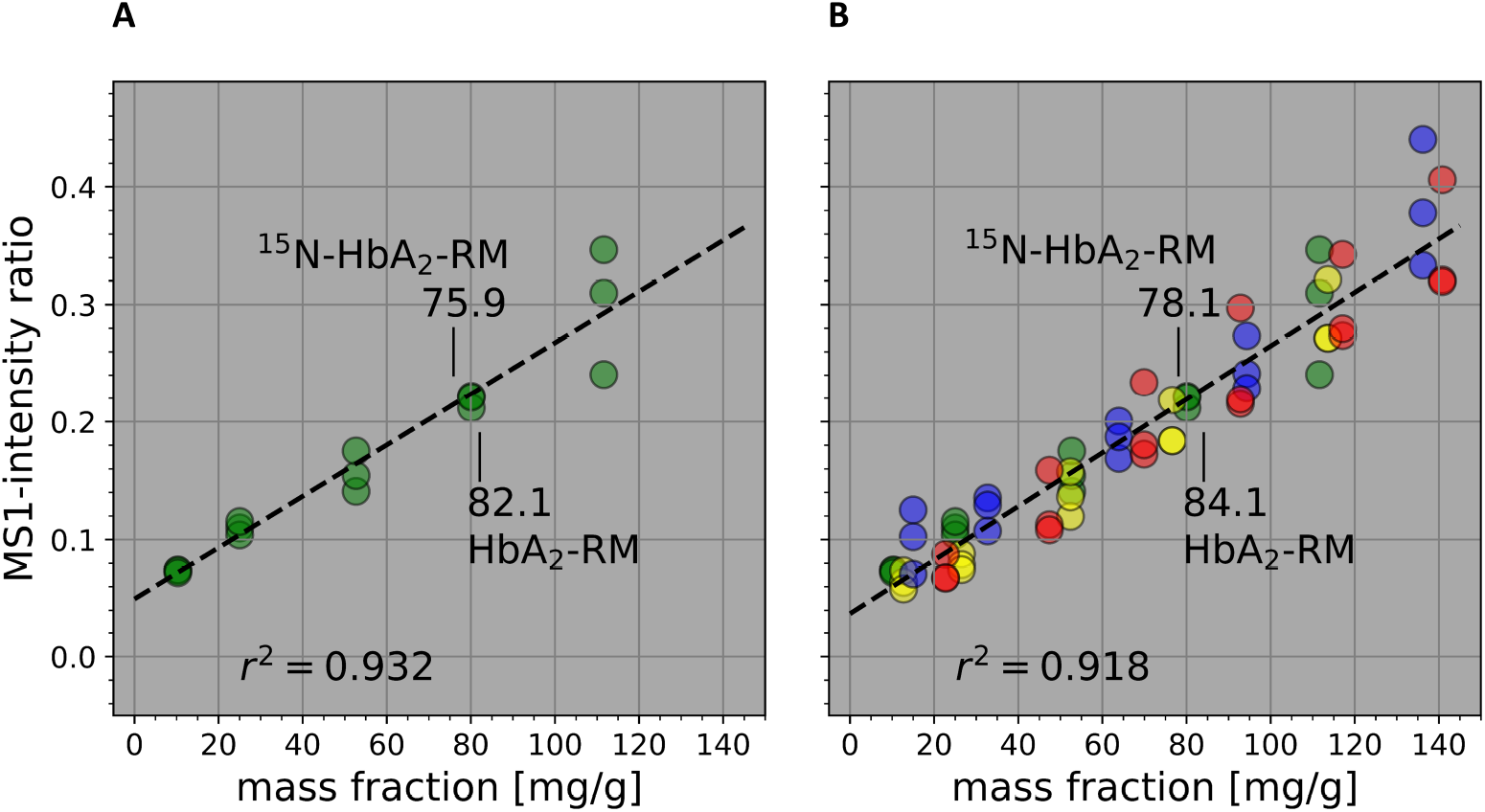
Calibrating MS1-intensity ratios (proteome ÷ HbA_2_) vs. mass fractions of HCPs-impurity in HbA_2_. (A) Calibration using *E.coli* -lysate, (B) Joint calibration using a set of different proteomes in addition to *E.coli : E.coli* -lysate (green), yeast (yellow), human K562-cell line (blue) and a mix of nine neat proteins (red). Dashed lines: linear regression fit. Annotations: results for the HbA_2_-RM and the U-^15^N-labeled RM using these calibrations.

### Representation of the sample RMs by the *E.coli* -lysate

*The* protein profiles in the RMs differ significantly from those in the lysate. This is illustrated by different aspects in Fig. 3. First, as expected, the set of identified *E.coli* -proteins in the RMs is significantly restricted relative to the cell lysate (Fig. 3A). Many of these are known to typically be copurified, if using IMAC for clean-up. Particularly, YfbG (P77398), YodA (P76344), GlmS (P17169) and ArgE (P23908) correspond to proteins previously reported in this context. ^13^ They make up a fraction of 60-70% (orange in Fig. 3B) in the RMs, while only about 4% in the lysate. The difference in protein profiles is further substantiated by a principal-components plot of results, as shown in Fig. 3C. Two series of samples of systematically changed *E.coli* -mass fractions are shown. The first one is simply the same data as was acquired with the *E.coli*-lysate calibration (green in Figs. 2A, 3A and C). The second one was generated by dilution of the labeled HbA_2_-RM (l53 proteins, blue in Fig. 3A and C). Unlike with most applications of principal components analysis (PCA), no data scaling was applied for the results in Fig. 3C. Object-scores (samples at different levels of mass-fraction) and feature-loadings (*E.coli* -proteins quantified) are jointly shown in the space of the first two components (PCl and PC2). In this kind of presentation, proximity of a protein (small black crosses) to a series (blue or green circles) corresponds with the involvement of that protein in the MSl-signal ratio for that series. At the same time, the distance from the origin quantitatively reflects the degree of this involvement. Visibly, the majority of proteins are significantly involved in just one of both materials. This particularly holds for the top abundant ones, such as YfbG. As an exception, on the other hand, YodA is one of the few markedly involved in both materials, though not to exactly the same extent.

**Figure 3:**
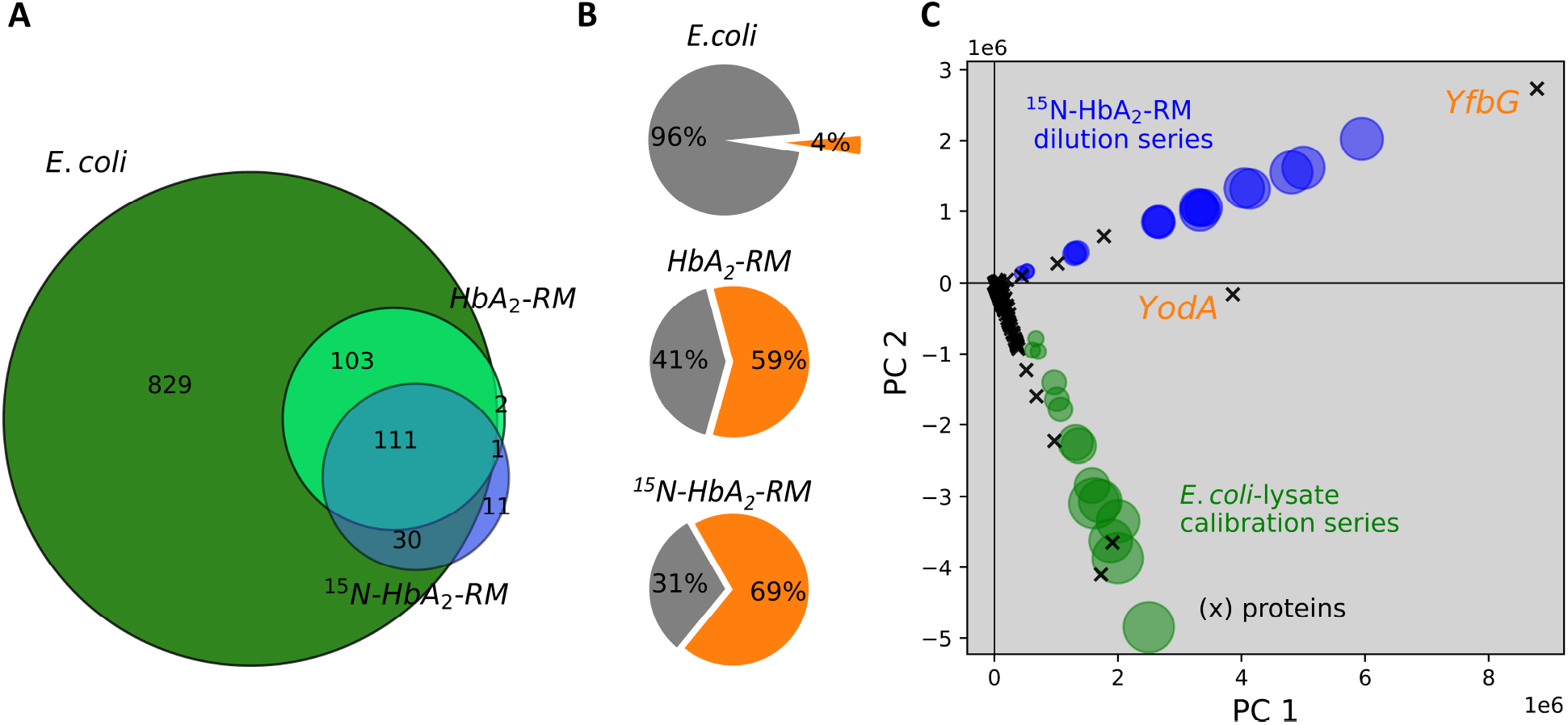
Comparison of the *E*.*coli*-protein profile in the lysate to the HbA_2_-RMs. (A) Identifications, (B) Relative amounts we found in these samples of frequently being co-purified^13^ *E.coli*-proteins (orange), (C) Interrelation of *E.coli*-protein fractions in *E.coli*-lysate (green circles) and in labeled HbA_2_-RM (blue) with MSl-intensities of individual proteins (x), illustrated by a principal-components plot. The series of protein fractions for the lysate is based on the same data source as the calibration data shown in Fig. 2A and 2B (green subset), whereas the HbA_2_-RM-series was obtained by dilution of the labeled HbA_2_ material. Areas of the circles are in proportion with the fractions in the pertaining samples. The plot is a projection of the data on the plane of the first two principal components, PC 1 and PC 2, of the joint dataset (lysate plus labeled HbA_2_-RM). Variance coverage: 61.6% (PC 1) and 36.1% (PC 2).

The demonstrated difference in protein profiles obviates prediction of impurity in an unknown material by linear regression models using as inputs individual proteins from another material (as e.g. *E.coli*). As previously mentioned, LFQ works around the problem by integrating signals over all proteins for *E.coli* on one hand in relation to HbA_2_, on the other side for mapping the mass fraction. This notion, indeed, has been consensus in the literature long since, ^21-23^ but still was taken to the test for the present purpose. To this, additional proteomes, beyond *E.coli*-lysate, were used for calibration, while otherwise subjected to the same workflow as before. These proteomes were: yeast, K562 and a mix of nine neat proteins.

The eventual reduction to the simple protein mix was on purpose to provide an artificial proteome with a minimum number of components. Results of the respective calibration runs are plotted in Fig. 2B HCPs-mass fractions obtained for the two HbA_2_-RMs by application of these calibrations are annotated in Fig. 4A and B. Apparently, there is good agreement as a whole between the individual plots. This supports the assumption that, the individual linear calibration models (per proteome) are samples from a common statistical population. This in turn suggests that calibration based on the *E.coli* -lysate (Fig. 2A) should essentially be valid for predicting HCPs-fractions in the RMs, too. Finally, pooling all individual calibrations into a common one is possible, as shown in the top trace (black) in Fig. 4, which may enhance the statistical robustness of value-assignments.

**Figure 4:**
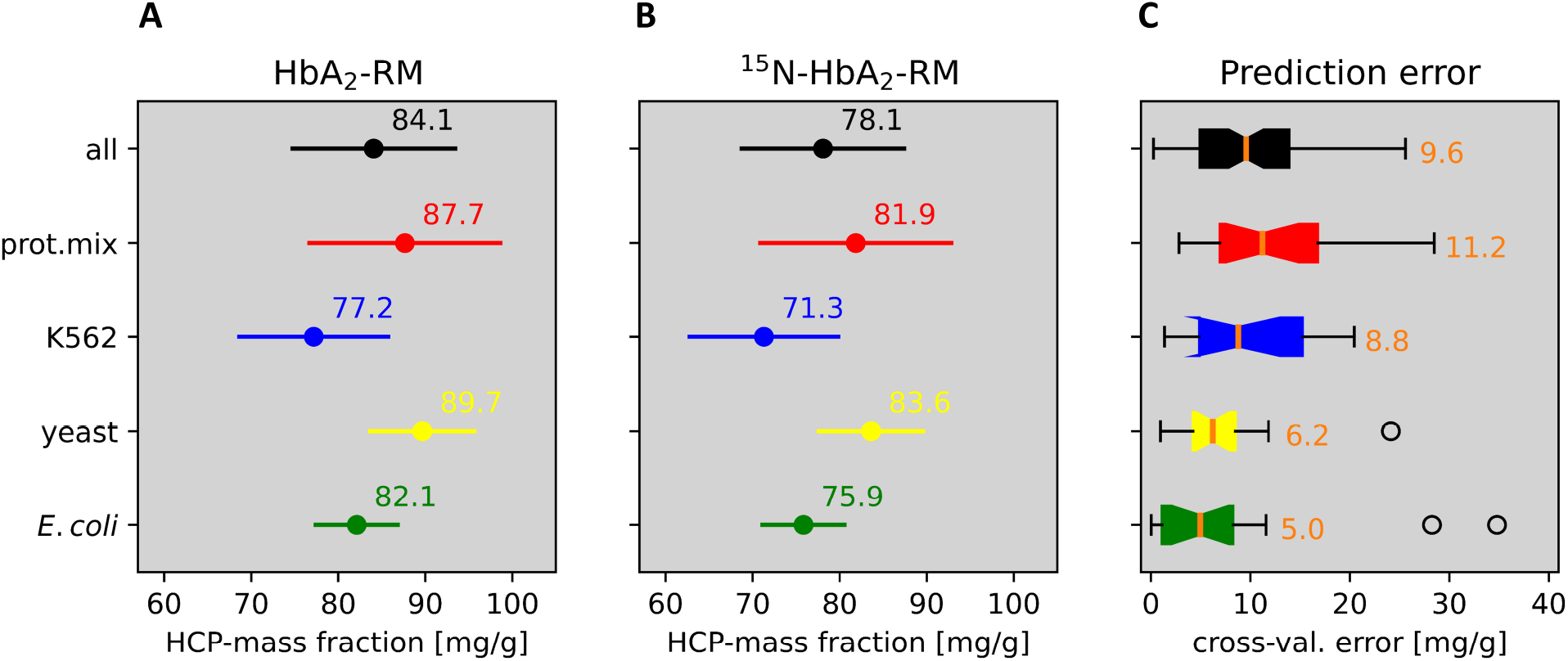
HCPs-fractions determined by LFQ for both HbA2-materials, and degree of equivalence between different calibrator-sources: *E.coli*-lysate (green), yeast (yellow), human K562 cell line (blue), protein mix (red) and all of these series merged into one (black). (A) Natural isotopic RM and (B) isotope labeled version; (C) distribution of prediction error obtained by leave-one-out crossvalidation of the calibration data; orange lines and numbers provide the medians. Circles: fliers. Error bars in (A) and (B) correspond to the medians in (C).

### Overall-uncertainty

Major sources of uncertainty are associated with: (i) mass fractions assigned to the calibrators (Fig. 1A) (ii) repeatability of sample preparation and measurement (Fig. 1B) and (iii) the fitness of the established linear models used for calibration (Fig. 1D). The calibrator-uncertainties *(i)* are likely to be dominated by value-assignment of mass fractions to the stock-solutions (HbA_2_, *E.coli*-lysate, yeast, K562 and protein mix). ID-MS based amino acid analysis (AAA) was employed for this, with an assigned uncertainty of 3.5% or less (see Experimental Section). For sample preparation and measurement *(ii)*, an estimate of 11.1% can be deduced from results of *n*= 6 repeated measurements of the labeled HbA_2_-RM. The contribution of the calibration function *(iii)* was finally evaluated by applying leave-one-out cross-validation to the calibration data. In detail, the calibration was established leaving out one of the calibrators, and its mass-fraction calculated then from the intensity-ratio using that calibration model. The difference to the known mass-fraction was used as an estimate of the prediction error. Fig. 4C shows the distribution of these errors having left out once, in this way, each of the calibrators, within each of the proteomes considered. Referring to the medians given in the Figure as estimates, the calibration-uncertainty would be about 11.2 mg/g, or less, which is about 13% relative to the results for the HCPs mass fractions in the two HbA_2_-RMs. Combining the three of the estimated uncertainty-components as a root sum of squares, provides an overall-uncertainty of 18% for a single measurement, in the case of the natural HbA_2_-RM. For the labeled HbA_2_-RM, on the other hand, with six repeats averaged, this would reduce to about 14%. Hence, values for HCPs-mass fractions can be assigned to the RMs of 84.1±15.0 mg/g (HbA_2_) and 78.1± 10.9 mg/g (^15^N-HbA_2_).

### Result for the *ultra-purified Hb*A_2_-material

To demonstrate scalability of the method, a separate calibration was performed in the range of 0.1−1.5 mg/g fraction of protein mix. Based on duplicate measurements of calibrators at 0.15, 0.3, 0.43, 0.71, 1.08, and 1.42 mg/g, a linear fit was obtained at a coefficient of determination of *r*^2^= 0.964, comparable to the previous broader-range calibrations (as e.g. 0.932 with *E.coli, cf*. Fig.2A). The calibration uncertainty by cross-validation was 0.086 mg/g, or 10%. Using the same uncertainty contributions as estimated before regarding stock solutions, as well as sample preparation and measurement, the result for this material is: 0.86±0.15 mg/g.

### Sample size and associated error

The previous results suggest that, calibration can in practice be performed with a small number of well characterized proteins quite as good as with complex biological materials. However, LFQ depends on the assumption that, the peptides captured by their MS1-intensities are drafts from the same population, for sample and calibrator, as regarding molar sensitivities. Consequently, on significantly reducing the number of peptide species involved, an associated sampling error will become apparent, thus increasing the overall measurement-uncertainty. In Fig. 5, results of a simulation are shown, attempting to estimate the size of that potential error. Calibration based on the protein mix was used and the deviation of obtained HCPs-mass fraction for the (unlabeled) HbA_2_-RM calculated, if reducing the number of proteins used for calibration down to *n* = 8−1 proteins, randomly selected from the nine. To generate a distribution of possible outcomes, 100 random drafts of this number of proteins were acquired at each level and the respective results for the HCPs-mass fraction were calculated. The data shown in Fig. 5 cannot exactly map reality, of course, since, even if using all of the nine proteins, the sampling error will be less than with just one, but cannot completely disappear at *n* = 9. As such, Fig. 5 does not. exactly reflect the ground truth, but it. should be close. Accepting this, the example suggests that, a number of five proteins may suffice on average to keep the sampling error at 15%, or less.

**Figure 5:**
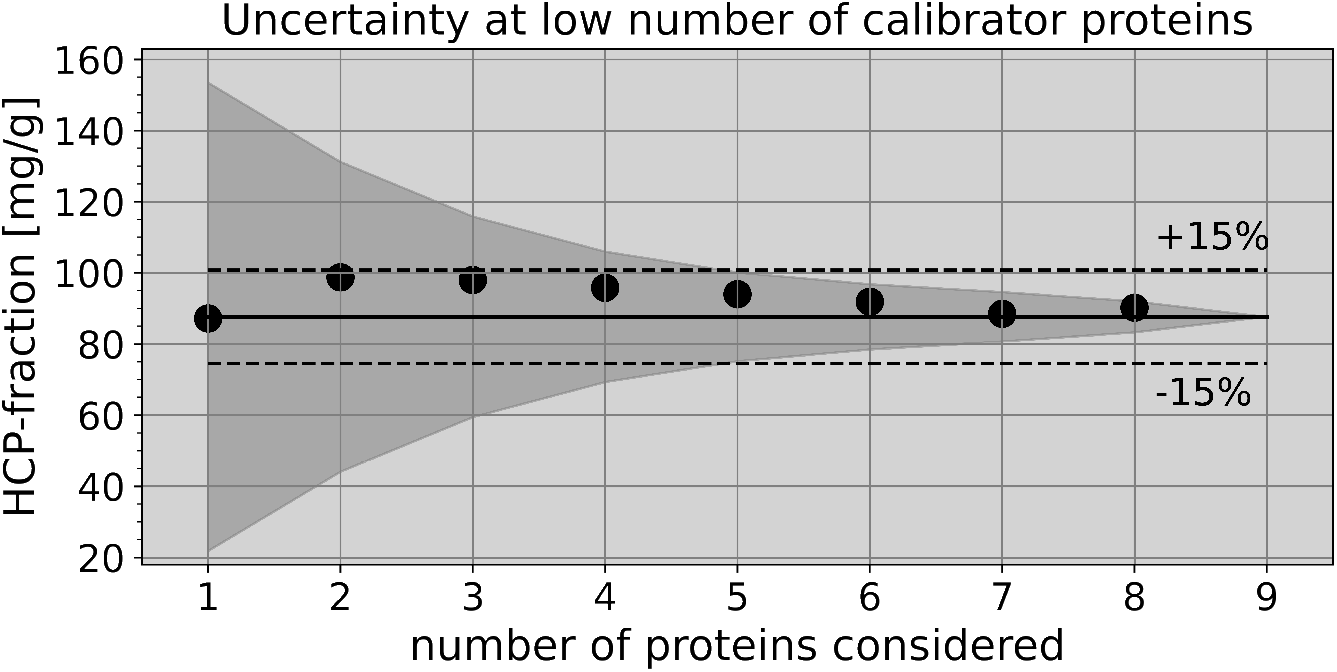
Estimating the sampling error caused by the finiteness of the number of proteins/peptides used for calibration, or present in the sample. The data shown are results for the HbA_2_-RM after stepwise reduction of the number of proteins included in calibration with the protein mix. Solid line: median obtained at *n* = 9 (87.7 mg/g), dashed: ±15%. Scatterpoints: median results after recalculating the calibration function using random drafts of *n* = 1−8 out of the originally nine proteins. Dark-grey area: corresponding standard deviations (shown here relative to the solid line, rather than to the medians).

### Top-N protein quantification strategy as an alternative

In the introduction we claimed an exhaustive one-by-one quantification of individual HCPs to be little practicable if the aim is to find the total amount (or fraction) of HCPs in cell-expressed proteins. Revision of the data shown in Fig. 3C indeed suggests the option of individually quantifying the top-5 *E.coli* proteins (YfbG and YodA for the most abundant ones) and taking the sum for an estimate. For the integrated MS1-signals, however, these five only account for some 88%, alltogether, for the ^15^N-HbA_2_RM compared to 69% for the natural HbA_2_-RM. \/here the impurity was of a yet more complex pattern, as with the *E.coli*-lysate shown in Fig. 3C, for instance, the result was even worse, with only 25% being accounted for by the five most abundant proteins in this example. Conceding that the requirement in relative precision may be relaxed to a certain degree for quantifying minor components like HCPs, when compared to the target protein, a remaining uncertainty of 12−75% of non-captured proteins is probably not satisfactory. A further argument in favor of LFQ it is that, the one-by-one approach is likely to be more expensive, compared to a series of simple shotgun-experiments, as required in LFQ.

## CONCLUSIONS

LFQ is applicable to the quantification of host-cell derived impurity in bioengineered proteins. Calibrating the integrated MS1-intensity for all HCPs against the same quantity obtained for samples of known mass-fractions is a straightforward solution to the problem of quantitatively capturing a composite set of individual proteins ultimately to be expressed as a gross-measurand. Viability of this proceeding is not hampered by the fact that the profile and identities of HCPs do not normally coincide with those of the calibrator material. This opens up the option of using proteomes for calibration other than those suggested by the expression system. This commutability of materials means that simple mixtures of well characterized proteins are also viable candidates.

For the natural HbA_2_ RM we estimate 84.1±15.0 mg/g (HbA_2_), for the isotope labeled HbA_2_-RM 78.1± 10.9 mg/g and for the purified (natural) HbA_2_-RM 0.86±0.15 mg/g. This translates to 916±15 mg/g, 922± 11 mg/g and 999.1±0.15 mg/g, respectively, fractions of HbA_2_ in the materials. The latter provide the correction factors to be applied to a quantitative result, if using these materials as reference. For the first two (IMAC purified) materials, an uncertainty result of ≈ 1−2% contributed to the overall budget for the analytical result, while ≈ 0.2% was obtained for the *ultra*-purified RM. Considering the expense in terms of material loss, and assuming a target of 5% uncertainty as acceptable for the protein as measurand in a biological sample, immediate use of the IMAC-purified material would have been optimal when compared to the efforts required for further purification.

Although discussed here in the context of value-assignment to RMs to be used as primary calibrators with protein quantification, LFQ is increasingly also being used in areas, such as process optimization and quality control of pharmaceutical products. ^24−28^ Typically in most of these applications, it is about quantification of individual proteins, rather than aiming at a mass fraction as a whole for HCPs. However, capturing a mass fraction of HCPs as a gross quantity, as discussed in this paper, or selectively for protein subclasses of particular interest, could gain importance in these industries for reasons of particular toxicity of such classes, or otherwise legal requirements.

